# Tracheal terminal cells of *Drosophila* are immune privileged to maintain their Foxo-dependent structural plasticity

**DOI:** 10.1101/2024.08.22.609264

**Authors:** Judith Bossen, Larissa Fritz, Reshmi Raveendran, Leizhi Shi, Jingjing He, Thomas Roeder

## Abstract

Respiratory organs must balance their primary function of gas exchange with the constant threat of inhaled pathogens. In the *Drosophila* tracheal system, gas exchange occurs at the tracheal terminal cells (TTCs), the functional equivalents of mammalian alveoli. While bacterial infection triggers a robust innate immune response throughout the broader airway epithelium, we reveal that TTCs are uniquely exempt from this reaction. Mechanistically, TTCs lack expression of the membrane-associated peptidoglycan recognition receptor PGRP-LC. This absence protects these highly susceptible cells from Immune deficiency (Imd) pathway activation and subsequent JNK-mediated cell death, establishing TTCs as a distinct, immune-privileged niche. Ectopic immune activation via targeted *PGRP-LCx* overexpression in TTCs caused a severe reduction in branching, cellular damage, and ultimately cell death, phenotypes that were fully rescued by the depletion of AP-1 or *foxo*. Because both structural plasticity (in response to nutritional cues and hypoxia) and innate immune responses strictly require the transcription factor FoxO, we demonstrate that potent immune signaling is fundamentally incompatible with dynamic TTC remodeling. Ultimately, the immune-privileged status of TTCs represents an essential evolutionary trade-off, restricting local inflammation to preserve *foxo*-dependent structural plasticity and vital respiratory function.

## INTRODUCTION

The tracheal system of *Drosophila* comprises three different parts: the main dorsal trunks, the smaller tracheal branches, and the terminal cells (Whitten 1957; Uv et al. 2003). The tracheal terminal cells (TTCs) form the most distal part of the respiratory system. They are the site at which gas exchange takes place. TTCs fulfill an essential role by supplying all body organs with oxygen. To fulfill their task, even under ever-changing conditions, these cells possess a high degree of structural plasticity. For example, TTCs can sprout in response to local hypoxia (Jarecki et al. 1999), or in response to differences in nutrient availability (Centanin et al. 2010); this process is analogous to angiogenesis in mammals. Sprouting is dependent on growth factors, as well as the hypoxia-inducible factor (HIF)-α homolog sima, again analogous to endothelial cells during angiogenesis in the mammalian vascular system (Jarecki et al. 1999; Centanin et al. 2008; Horowitz and Simons 2008).

The primary role of the proximal tracheal tubes is to conduct air to more distal parts. The epithelial cells lining these tubes can mount an effective immune response to infections (Wagner et al. 2008; Faisal et al. 2014). This immune response is driven mainly by expression of several antimicrobial peptides (AMP) (Tzou et al. 2000; Akhouayri et al. 2011). One out of the two major immune pathways operative in *Drosophila*, the immune deficiency (Imd) pathway, is solely responsible for responses in the larval airway epithelium; this is because the Toll pathway is not functional in these cells (Wagner et al. 2008) Klicken oder tippen Sie hier, um Text einzugeben.. The Imd signaling pathway is homologous to the human tumor necrosis factor-α (TNF-α) pathway and converges on activation of NF-κB factors. Moreover, it is connected to c-Jun N-terminal kinase (JNK) signaling via transforming growth factor-β (TGF-β) activated kinase 1 (dTak1) (Silverman et al. 2003). Infection of the airway system induces an immune response by the tracheal epithelium, which includes expression of canonical Imd and JNK target genes (Tzou et al. 2000; Wagner et al. 2009). Chronic activation of tracheal immune signaling leads to marked structural changes in the epithelium (Wagner et al. 2021). This tissue remodeling is mediated by JNK and its downstream transcription factor, forkhead box sub-group O (foxo), which is a as a terminal target of the epithelial immune system. FoxOs are also central proteins of the insulin signaling pathway, which also plays an important role in the structural plasticity of TTCs (Linneweber et al. 2014; Wong et al. 2014; Burguete et al. 2019).

The present study shows that TTCs differ fundamentally from the rest of the tracheal epithelium. In the case of a natural infection of the tracheal system, these cells show only a negligible immune response. The very few TTCs that do show AMP expression are structurally impaired. We found that TTCs are immune privileged. The Imd pathway in TTCs is deactivated since the cells do not express the transmembrane peptidoglycan recognition protein (PGRP)-LCx. Chronic activation of the Imd signaling pathway in TTCs leads to JNK-dependent apoptosis. Therefore, we hypothesize that the immune privileged status of TTCs maintains the normal function of foxo, which acts as a regulator of structural plasticity.

## RESULTS

### Tracheal infection revealed that AMP expression in terminal cells is rare

Larvae carrying reporters for antimicrobial peptide genes were studied following infection with *Pectobacterium carotovorum* (formerly *Erwinia carotovora*) or under control conditions. These larvae express GFP under the control of promoters for different antimicrobial peptide (AMP) genes, which are known to be activated by a natural infection in different tissues (Tzou et al. 2000). GFP fluorescence was analyzed after 24 h, focusing on the TTCs (Fig. 1). Without infection AMP expression was rarely or never observed in the tracheal system (Fig. 1A-E). Expression in the TTCs was never observed. In infected animals, AMP expression was observed frequently in the tracheal system, but again, never in the TTCs (Fig. 1A’-E’). The emergence of AMP expression was quantified by counting the number of animals with fluorescence in the 3^rd^ tracheal metamere with focus on the dorsal part, especially in the dorsal branch/fusion cells (Fig. 1F; GFP+, light green) and fluorescence in the TTCs (GFP TTCs, dark green). In 5-33% of the larvae, fluorescence was detected in the dorsal branch (DB)/fusion cell (FC) region; this number increased after infection to 24-70 % (Fig. 1G). But infection could not trigger an antimicrobial peptide response in the TTCs. In addition to a response in the trachea, fluorescence was observed in various tissues, such as hemocytes and fat body, especially in the anterior part, indicating the animals’ general responsiveness to infection (Fig. S1A-D).

**Fig. 1:**
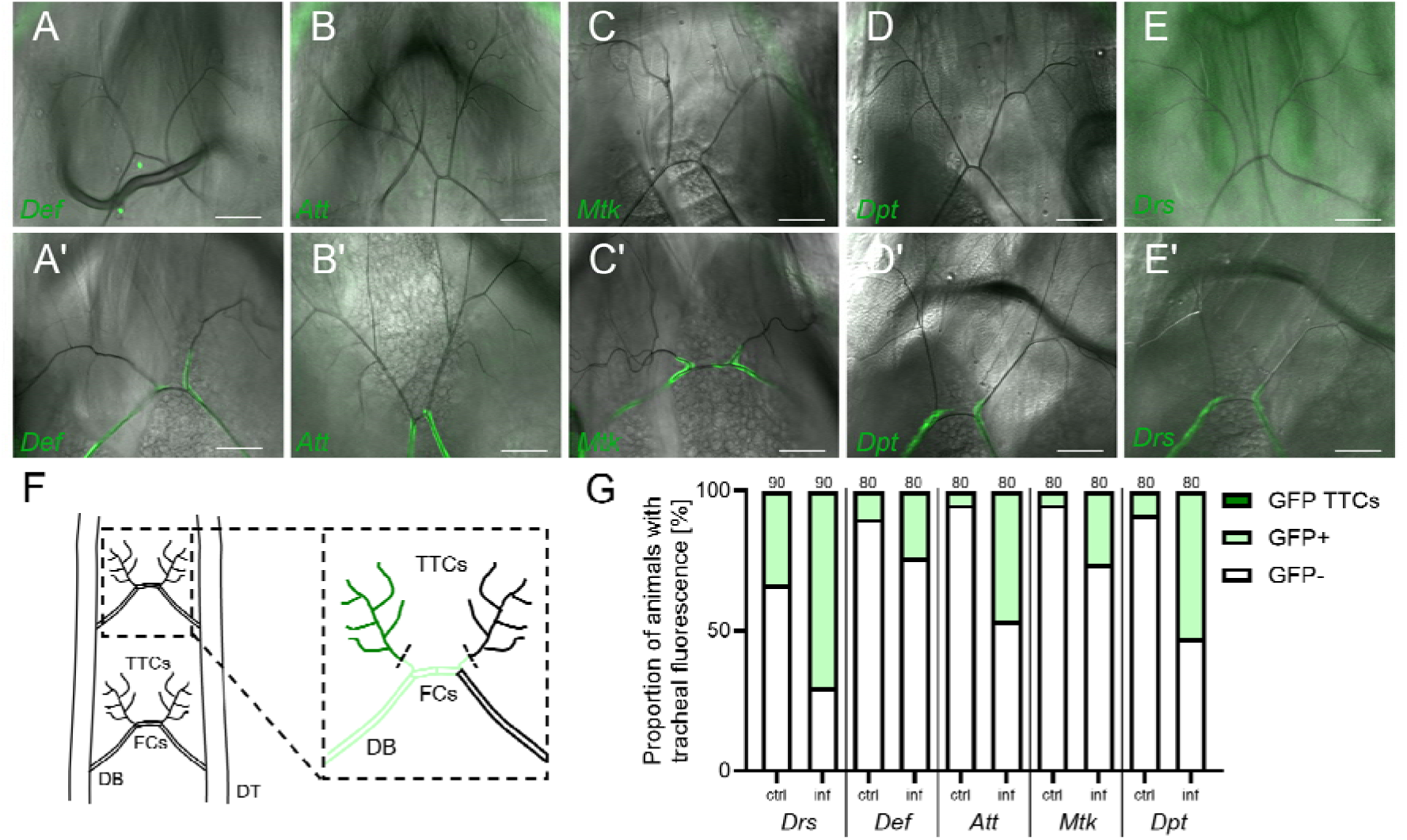
Terminal tracheal immune reaction to natural infection measured via AMP reporter response. GFP reporter larvae were infected with *P. carotovorum* for 24 h, and GFP fluorescence in the terminal structures of the tracheal system was monitored. The Images were taken in the DIC and GFP channels. Scale, 50 µm. (**A-E**) Dorsal tracheal structures of non-infected larvae. (**A’-E’**) Dorsal tracheal structures of infected larvae. (**F**) Dorsal view of the tracheal system, showing the dorsal trunks branching into the dorsal branch (DB), fusion cells (FC) and the dorsal TTCs. The quantification explanation is shown in magnification emphasized with colors. Presence of fluorescence in DB/FC counts as a GFP positive larva (light green, GFP+). Presence of fluorescence in the TTCs in GFP+ animals is indicated by dark green color (GFP TTCs). (**G**) Proportion of animals with tracheal fluorescence following the mentioned quantification procedure. Scale bar, 50 µm. Drs = Drosomycin, Def = Defensin, Att = Attacin, Mtk = Metchnikowin, Dpt = Diptericin.

For a more detailed analysis, we focused on the AMP Drosomycin, (*Drs*) which showed the strongest response to infection in the trachea. A total of 169 infected larvae were examined over all tracheal metameres. About 34% showed no GFP signal in the focus area, a finding consistent with previous observations (Akhouayri et al. 2011). The remaining larvae showed a GFP signal in the DB/FCs (57.4%, Fig. 2A, A’). The absence of a response in the TTCs was not limited to the dorsal TTCs but was also observed in TTCs supplying other body parts (Fig. 2B). However, 8.3% showed *Drs* expression in TTCs (Fig. 2C-G). Most of the few fluorescent TTCs showed clear structural differences from TTCs without a GFP signal (Fig. 2D, D’). Although some cells had a normal structure (Fig. 2E, E’), most had shortened branches (Fig. 2D, D’, F, F’), showed signs of melanization (Fig. 2G, arrow), or were no longer air-filled (Fig. 2G, arrowhead). These observations suggest an immune triggered response in the TTC itself that act negatively on these cells.

**Fig. 2:**
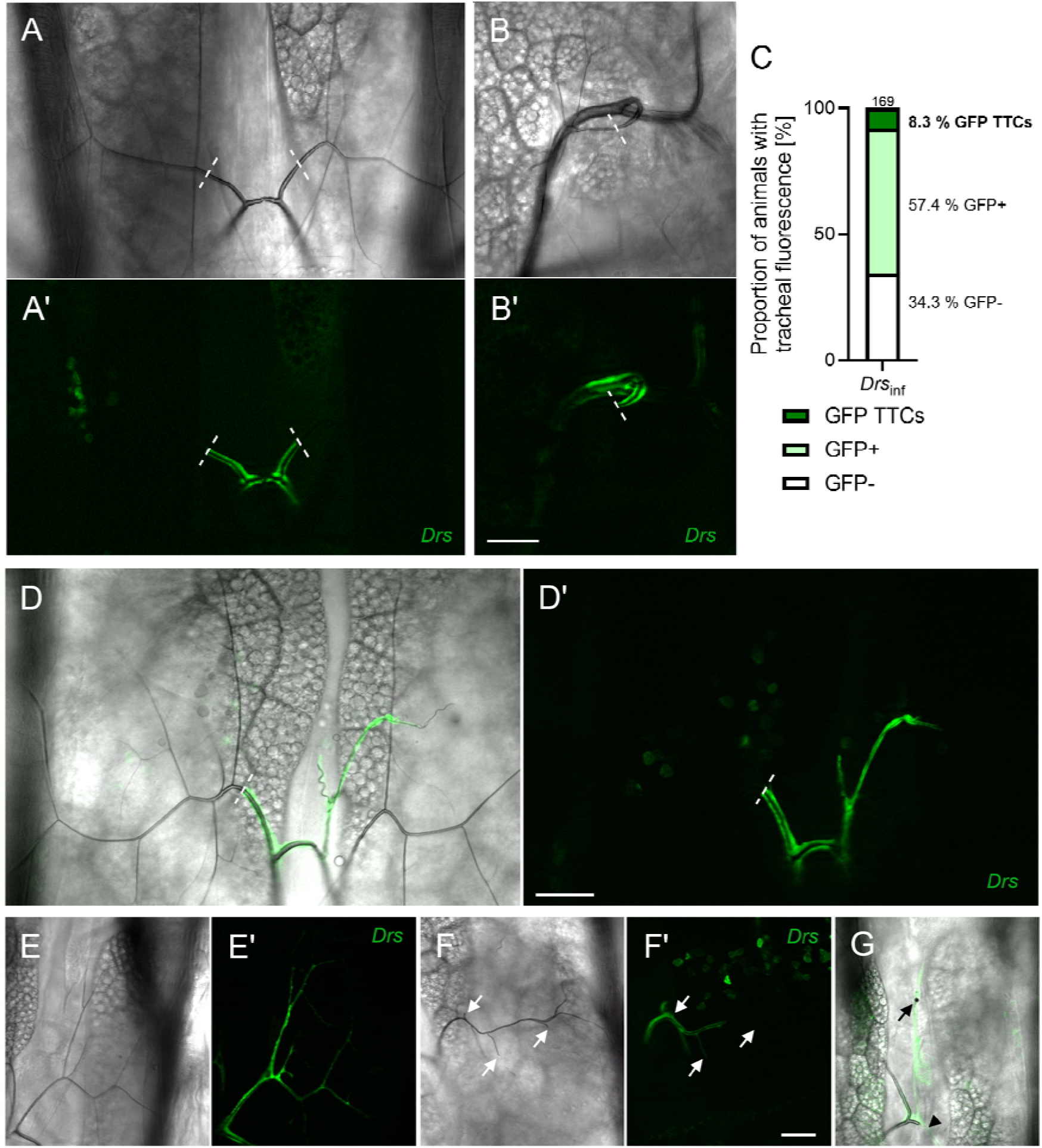
Tracheal terminal cells (TTCs) show very rare *Drs* expression upon natural infection. *Drs-GFP* larvae were infected with *P. carotovorum* for 24 h, and GFP fluorescence in the TTCs of the tracheal system was monitored. Images were taken in the DIC (A, C-G) and GFP channels (A’, C’-F’). (**A**) Dorsal TTCs without fluorescence. (**B**) Visceral TTCs without fluorescence. (**C**) Percentage of larvae showing GFP fluorescence in the DB and TTCs. (**D**–**G**) TTCs with expression of *Drs-GFP*. White arrows indicate shortened TTC branches (G). The black arrows mark a melanization site and the arrowhead marks a translucent branch without air filling (G). Dashed lines represent the proximal end of the TTCs. Scale bar, 50 µm.

### Tracheal terminal cells do not express the Imd-pathway receptor PGRP-LCx

We wondered whether constitutive immune activation within the tracheal system would exclude TTCs, as expected from the results of the infection experiments. To ensure that all parts of the tracheal epithelium, including TTCs, were exposed to this activating stimulus, we ectopically expressed the secreted (and intracellular) Imd pathway receptor PGRP-LE using the *ppk4-Gal4* driver. AMP expression (a readout of the immune response) was visualized by simultaneous expression of GFP-tagged *Drs* (Fig. 3). A strong immune response was observed in the tracheal tissue, from the dorsal trunks down to the smaller branches (Fig. 3A–C). While expression of *Drs* was consistent throughout most parts of the tracheal system, it was absent from the most distal parts (Fig. 3B–E). Closer observation of the TTCs revealed a distinct breakpoint in the GFP signal at the proximal end of the TTCs. This was true for TTCs attached to the outer cuticle (Fig. 3D) as well as for cells attached to the intestine (Fig. 3E). Activation of the Imd pathway may not be fully functional in these cells. We infected the animals and quantified the corresponding response in the 3^rd^ tracheal metamere. We observed an increased response upon infection and a few fluorescence events in the TTCs in 3% of the animals (Fig. 3F, G).

**Fig. 3:**
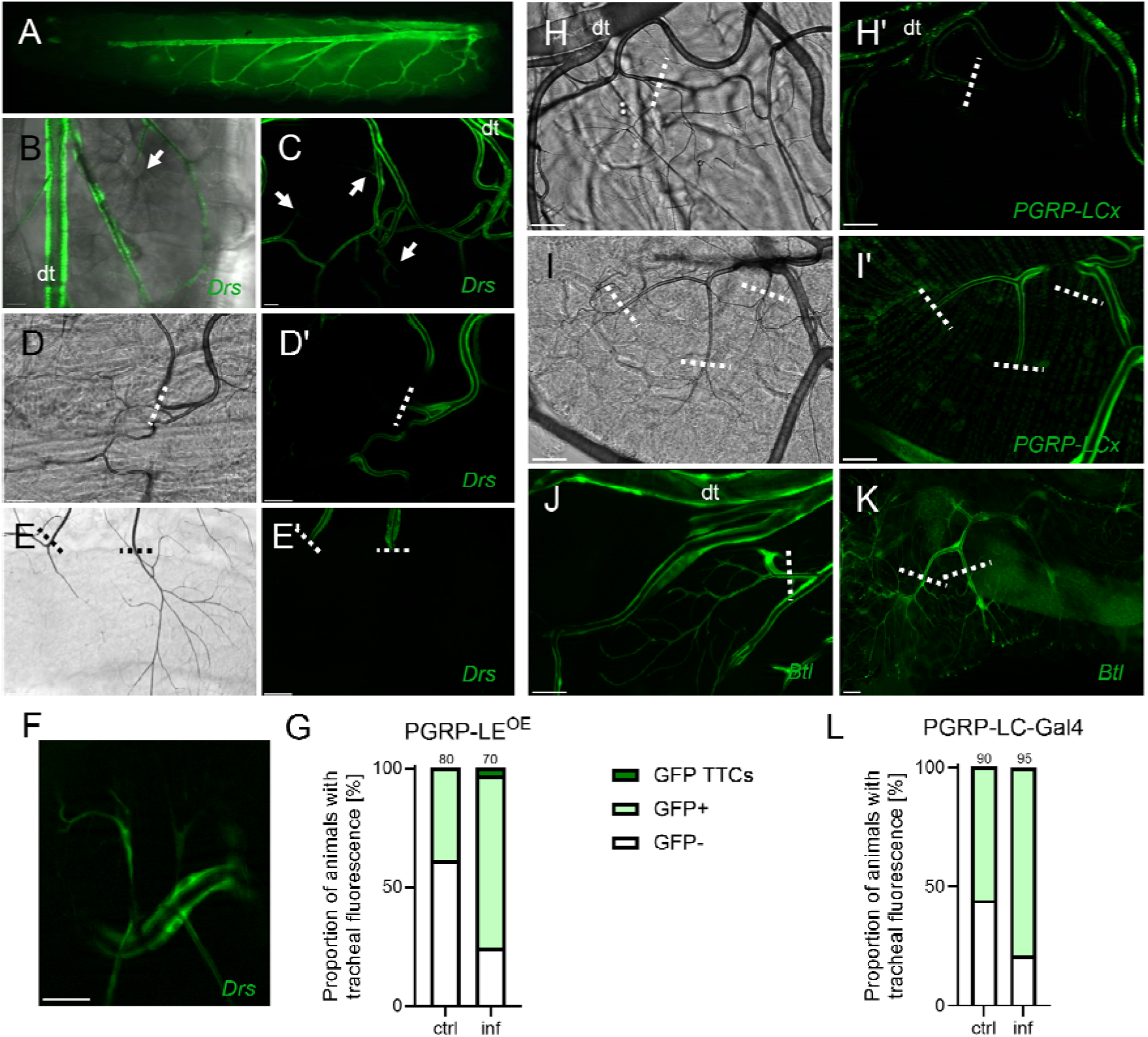
TTCs do not express the Imd receptor PGRP-LCx. **(A–D)** The secreted Imd pathway receptor *PGRP-LE* is expressed in the main parts of the tracheal system (*ppk4>PGRP-LE (GFP-Drs)*). The arrows indicate TTCs not expressing GFP. The dashed lines represent the proximal end of the TTC. **(A–C)** An activated immune response in the larvae is visualized by expression of GFP-tagged *Drosomycin* (*Drs*). **(D, E)** Detailed TTCs were observed in fillet preparations (D) and in the dissected intestine (E) in both the DIC (D, E) and GFP channels (D’, E’). (**F**) Example TTC of infected PGRP-LE animals with fluorescence. (**G**) Percentage of PGRP-LE^OE^ animals with tracheal GFP expression. **(H, I)** Expression of GFP under the control of a *PGRP-LCx* promoter (*PGRP-LCx-Gal4 > UAS-GFP*) revealed a lack of promoter activity, and expression of GFP, in TTCs on the cuticle (H, H’) and intestine (I, I’), which is in contrast to the rest of the tracheal system. **(J, K)** The TTCs in the tracheal system are visualized by GFP expression in the cuticle (J) and intestine (K) of wild-type larvae (*btl-Gal4; UAS-GFP*). (**L**) Percentage of *PGRP-LCx* animals with tracheal GFP expression. Scale bar, 50 µm. Dashed lines represent the proximal end of the TTCs. Dt = dorsal trunk.

To activate the Imd pathway successfully, secreted PGRP-LE has to bind to transmembrane PGRP-LC, meaning that only cells expressing PGRP-LC can be activated (Takehana et al. 2002; Takehana et al. 2004). Therefore, to investigate PGRP-LC expression in the tracheal system, specifically in TTCs, we expressed *Gal4* controlled by the *PGRP-LC* promoter to drive concurrent GFP expression (Fig. 3H, I). While GFP was visible throughout the entire tracheal system, it was absent from all TTCs associated with the cuticle (Fig. 3H, H’) and intestine (Fig. 3I, I’). To demonstrate that GFP expression can be visualized in TTCs, and that the lack of a signal was not an artifact, expression of GFP in TTCs was driven by the tracheal driver *btl-Gal4* (*btl-Gal4; UAS-GFP*) (Fig. 3J, K). TTCs on the dissected cuticle (Fig. 3J) and on the intestine (Fig. 3K) expressed GFP. Similarly, as before, we infected the *PGRP-LCx-P-Gal4* larvae and observed an increased number of animals with fluorescence in the 3^rd^ metamere DB/FCs but no animals with fluorescence in the TTCs (Fig. 3L).

Thus, the data indicate that lack of expression of the Imd-pathway receptor PGRP-LC by TTCs is the reason for the difference between TTCs and cells within the rest of the tracheal system. The observation that only 3 % of the PGRP-LCx expressing animals show *Drs* expression in the TTCs after infection (Fig. 3F, G) opens up alternative modes of induction including potential stress responses mediated through e.g., Foxo activation.

### Ectopic activation of PGRP-LCx in the tracheal system leads to the death of TTCs

To further investigate the effects of PGRP-LCx-mediated immune responses on the tracheal system, we expressed *PGRP-LCx* exclusively in TTCs (*PGRP-LCx^OE^*; Fig. 4). We found that Imd-mediated immune activation was restricted to these cells, as indicated by concurrent expression of GFP. Compared with the widely branched and diversified TTCs in control samples (Fig. 3A), *PGRP-LCx*-expressing TTCs showed a striking phenotype characterized by smaller TTCs with shortened terminal branches (Fig. 4A, B). Measurement of TTC branches in *PGRP-LCx*-expressing larvae revealed a significant reduction in the number and length of branches compared with the controls (Fig. 4C, D). While cells in the control had about 14 branches, with a total length of about 1550 µm, TTCs in PGRP-LCx^OE^ (III) insects had seven branches at the most, and the length did not exceed 514 µm. This means that the branched surface of TTCs expressing *PGRP-LCx* was reduced to less than one-third of that in controls. Additionally, we tested another PGRP-LCx expressing line (PGRP-LCx^OE^ (II), also used later for mechanistic evaluation). We could see a less severe but still prominent TTC-phenotype (Fig. 4C, D). To prove the less severe phenotype, we drove PGRP-LCx^OE^ (III) and (II) in the dorsal trunks of the trachea and compared the tracheal thickness (Fig. S2A-D). The PGRP-LC-dependent epithelial thickness phenotype was shown in a previous study (Wagner et al. 2021). PGRP-LCx^OE^ (III) produced a significantly increased thickening phenotype compared to PGRP-LCx^OE^ (II) (Fig. S2A-D).

**Fig. 4:**
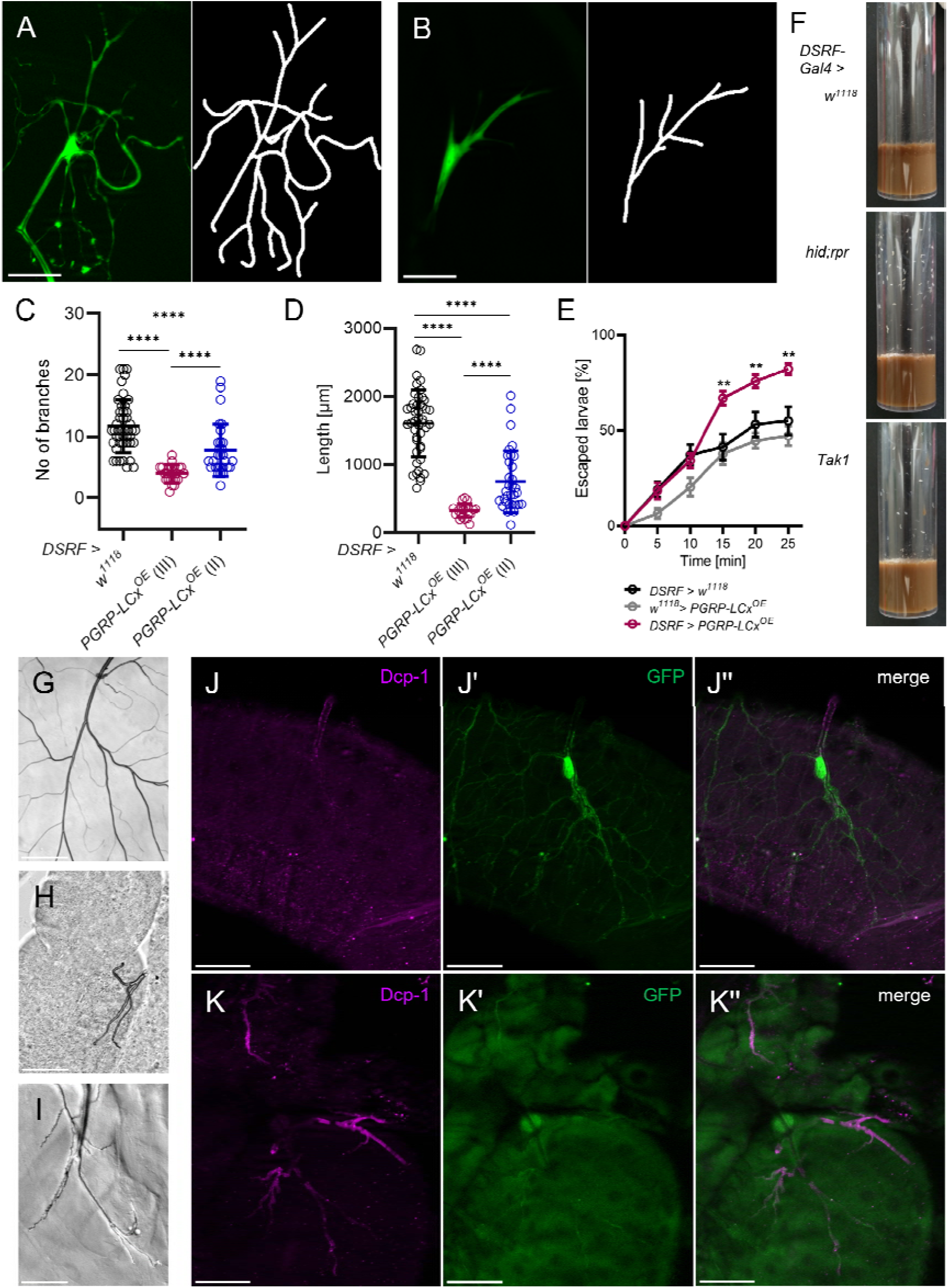
Expression of PGRP-LCx by TTCs leads to size reduction and loss of functionality. **(A, B)** Dorsal TTCs in the control (A) and *DSRF*-driven overexpression of *PGRP-LCx* in TTCs (B). **(C, D)** Measurement and quantification of the number (C) and length (D) of branches (n=22–45). Data are presented as the mean ± SD. **(E)** The hypoxia sensitivity assay was conducted with control and *PGRP-LCx*-expressing 3^rd^ instar larvae under hypoxic conditions (2–3 % O_2_, n = 11–14). Data are presented as the means ± SEM. Statistical significance was tested using Mann-Whitney-U test. * p < 0.05, ** p < 0.01, *** p < 0.001, **** p < 0.0001. (**F**) Culture vials containing control, hid;rpr, and Tak1 larvae (*DSRF > hid;rpr/Tak1*) at 4 days post-oviposition. **(G, H)** Transmission light microscopy of dissected intestines from 3^rd^ instar larvae (F, G) or 2^nd^ instar larvae (H) with the connected TTCs of control (G) showing expression of *hid;rpr* (H) or *PGRP-LCx* (I). **(J, K)** Dissected intestines from control larvae (J) or larvae expressing *PGRP-LCx* in TTCs (K) were stained with an antibody specific for cleaved *Drosophila* Dcp-1 (purple). (**I’**, **J’**) and then counterstained to detect GFP (green). (**I’’**, **J’’**) Merged channels. Scale bars, 50 μm.

To investigate whether this cellular phenotype has an impact on physiological oxygen supply, we exposed larvae to low oxygen levels (2–3 %). Leaving the lawn under hypoxic conditions is the normal response of larvae, and this response can be monitored over time. A significantly higher percentage of larvae with TTCs overexpressing *PGRP-LCx* showed the escape phenotype after 15–25 minutes of exposure (Fig. 4E, S4). Activation of the Imd pathway by ectopic expression of *PGRP-LCx* clearly impairs TTCs, not only with respect to size reduction, but also to the functionality of these cells (i.e., the enhanced response to hypoxic conditions). To elucidate whether this size reduction is part of an apoptotic cell program, we analyzed expression of two apoptotic factors: head involution defective (hid) and reaper (rpr) (White et al. 1994; Grether et al. 1995). We targeted expression of these factors to TTCs using the *DSRF*-Gal4 driver line. Larvae expressing *hid; rpr* showed a strong response to hypoxia by leaving the lawn at the very early larval stages; these larvae did not survive (Fig. 4F). Since PGRP LCx-expressing larvae survived and did not show this extreme phenotype, we examined Tak1 (a downstream component of the Imd pathway). Expression of *Tak1* in TTCs also resulted in early larval death, similar to expression of *hid; rpr* (Fig. 4F). Dissection of *hid; rpr*-expressing insects revealed a complete absence of TTCs, with only rudimentary cell remnants connected to the intestine (Fig. 3G, H). By contrast, overexpression of *PGRP-LCx* in TTCs resulted in similar cellular alterations, although the cells remained alive. Nevertheless, the overall phenotype was strikingly similar to that of apoptotic terminal cells (Fig. 4I).

*Drosophila* death caspase-1 (Dcp-1) is orthologous to mammalian Caspase-3, and is an established marker of apoptosis (*21*). Immunohistochemical analysis using an anti-Dcp-1 antibody (Fig. 4J, K) revealed a lack of Dcp-1 in the control; however, strong GFP staining of TTCs was visible (Fig. 4J-J’’). TTCs expressing *PGRP-LCx* were positive for cleaved Dcp-1, whereas counterstaining for GFP revealed reduced expression (Fig. 4K-K’’). These data could indicate that TTCs are undergoing apoptosis in consequence of PGRP-LCx overexpression. To complete our results, we stained trachea with PGRP-LCx expression in the tracheal epithelium and saw an increase of cleaved Dcp-1 in these tracheae as well (Fig. S3).

We expressed PGRP-LE in the trachea for only 24 h to induce a mild activation of the pathway in TTCs and epithelial cells (using *btl-Gal4; tubGal80ts*) and compared hypoxia response under control and infected conditions (Fig. S4). While we saw the same increase in the escape response of PGRP-LE larvae under control conditions, the response was even more enhanced for the infected larvae.

### PGRP-LCx-induced terminal cell death is mediated by JNK signaling

The *Drosophila* Imd signaling pathway, which classically leads to activation of the Nf-κB factor Relish, is homologous to the TNF-α signaling pathway in mammals. Like the TNF-α pathway, the Imd pathway branches out to the JNK signaling pathway; the branch point is Tak1 (Fig. 5A). To test whether the Nf-κB or the JNK branch is necessary to translate PGRP-LCx activation in TTCs into triggering of apoptosis, we expressed these downstream components in flies, which were then phenotyped. The number and length of the terminal branches were measured and compared with those in control TTCs, and with the phenotype produced by *PGRP-LCx* expression (Fig. 5B–F). Ectopic expression of an activated *Relish* allele had no effect on the TTC branching phenotype (Fig. 5B, E, F). Expression of a constitutively active form of dJNKK, *hemipterous* (*hep^CA^*), resulted in a phenotype like that caused by expression of the upstream receptor PGRP-LCx (Fig. 5C, E, F). Branching of TTCs was reduced significantly and did not differ significantly from that of TTCs with *PGRP-LCx* expression. Overexpression of the wild-type JNK, *basket* (*bsk^OE^*), also resulted in reduced branching when compared with the control (Fig. 5D–F). These results suggest that JNK activation is responsible for the TTC phenotype. To further support this hypothesis, the cells were stained for the Nf-κB factor Relish (Rel) and for the phosphorylated JNK (pJNK) basket (Fig. 5G, H).

**Fig. 5:**
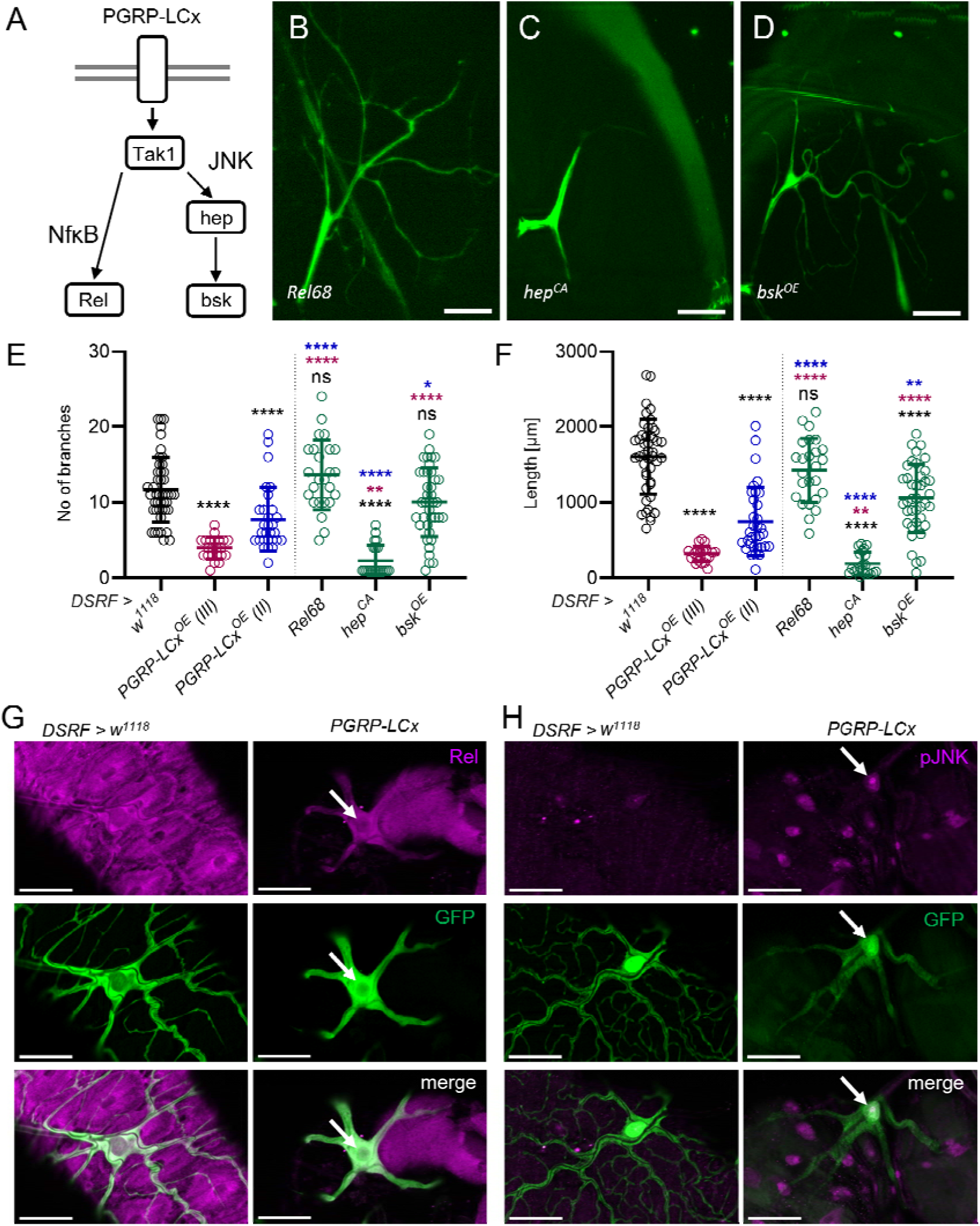
JNK signaling is associated with impaired TTC branching. **(A)** Schematic showing how the PGRP-LC-activated Imd signaling pathway is subdivided into the NF-κB (Relish) and JNK (hep = JNKK, bsk = JNK) pathways. **(B–D)** Relish (B, *Rel68*), as well as constitutively active hep (C, *hep^CA^*) and bsk (D, *bsk^OE^*), were expressed in TTCs (*DSRF >*). **(E, F)** Measurement and quantification of the number (E) and length (F) of branches in control (*w^1118^*) versus *PGRP-LCx*-expressing TTCs (n=22–45). Data are presented as the mean ± SD. Statistical significance was evaluated using Mann-Whitney-U test, ** p < 0.01, **** p < 0.0001, ns = not significant. **(G, H)** Dissected intestines from control (*DSRF > w^1118^*) and *PGRP-LCx*-expressing flies (*DSRF > PGRP-LCx*), in which the TTCs were stained to detect Relish (Rel, G) and pJNK (purple, H). TTCs were counterstained with GFP (green). Arrows mark the TTC nucleus. Merged channels are shown. Scale bars, 50 µm.

Staining of the transcription factor Relish was observed throughout intestinal tissue, as well as in TTCs, from controls, and in TTCs expressing *PGRP-LCx*; however, we observed no translocation of Relish to the nucleus (Fig. 5G, arrow). In contrast to controls, *PGRP-LCx*-expressing cells showed distinct nuclear staining by an anti-pJNK antibody (Fig. 5H, arrow). Moreover, nuclear pJNK was also observed in the nuclei of subjacent intestinal cells. This suggests that intestinal cells experience some additional JNK-mediated stress.

### JNK-mediated TTC damage can be rescued by AP-1 and foxo depletion

The *Drosophila* JNK signaling pathway induces apoptosis (Adachi-Yamada et al. 1999; Takatsu et al. 2000). The canonical transcription factors (TFs) AP-1, foxo and Ets21C act downstream of Tak1 and JNK to initiate transcription of target genes (Fig. 6A). Activation of each of these factors can lead to expression of apoptotic genes (Hafezi et al. 1997; Lehmann et al. 2002; Luo et al. 2007; Mundorf et al. 2019). Klicken oder tippen Sie hier, um Text einzugeben.To evaluate whether one of these TFs is responsible for the *PGRP-LCx^OE^*-induced phenotype, we silenced each of them. This experimental approach required the use of different controls with the *PGRP-LCx^OE^* cassette located on different chromosomes. Beside the PGRP-LCx^OE^ (III) on the 3^rd^ chromosome, we used PGRP-LCx^OE^ (II) which produces a less severe phenotype and is located on the 2^nd^ chromosome (Fig. 6F, G; red and blue). First, we used a dominant-negative form of *Tak1* (*Tak1^DN^*) and *bsk* (*bsk^DN^*) together with *PGRP-LCx^OE^* to confirm that PGRL-LC-induced apoptosis of TTCs is mediated by JNK. Concurrent expression of *PGRP-LCx^OE^* with either *Tak1^DN^* or *bsk^DN^* rescued the apoptotic phenotype (Fig. 6B, F, G). Fos and Jun proteins form homo- or heterodimers that act as AP-1 transcription factors, also a canonical target of the JNK pathway. To target AP-1, we focused on the *Drosophila* Fos ortholog kayak (kay). We co-expressed a dominant-negative *kay* allele (*kay^DN^*) together with *PGRP-LCx^OE^*in TTCs. Although *kay^DN^* rescued the *PGRP-LCx^OE^*phenotype with respect to the number and length of branches, it was not complete (Fig. 6C, F, G; red). Co-expression of *foxo^RNAi^* rescued the *PGRP-LCx^OE^* phenotype to the same level as *kay^DN^* (Fig. 6D, F, G; red). RNAi of *Ets21C* failed to rescue the *PGRP-LCx^OE^* phenotype (Fig. 6E, F, G; blue); rather, it worsened the phenotype with respect to the length of branches. Because depletion of the TFs foxo and kayak rescued the TTC phenotype induced by *PGRP-LCx* overexpression, we next tested whether upregulation of either induces a similar TTC phenotype (Fig. 6H–K). Overexpression of *foxo* led to a slightly reduction in the number and length of the branches (Fig. 6H–J), while overexpression of a combination of *kay* and *Jra*, which drives concurrent overexpression of both AP-1 components, led to a severe reduction in number and length of the branches, to levels below that induced by *PGRP-LCx* overexpression (Fig. 6H–J, K). Therefore, we asked whether foxo and AP-1 are present and active in TTCs. To do this, we used an AP-1 responsive TRE-RFP reporter line to detect AP-1 activity in TTCs (Chatterjee and Bohmann 2012). We observed AP-1 activity in wild-type TTCs, and upon overexpression of *PGRP-LCx* (Fig. 6L, M). When we used a *Gal4* line with a *foxo* promoter, we observed strong *foxo* promoter activity in all TTCs (Fig. 6N). Thus, both TFs appear to be expressed and functional in TTCs.

**Fig. 6:**
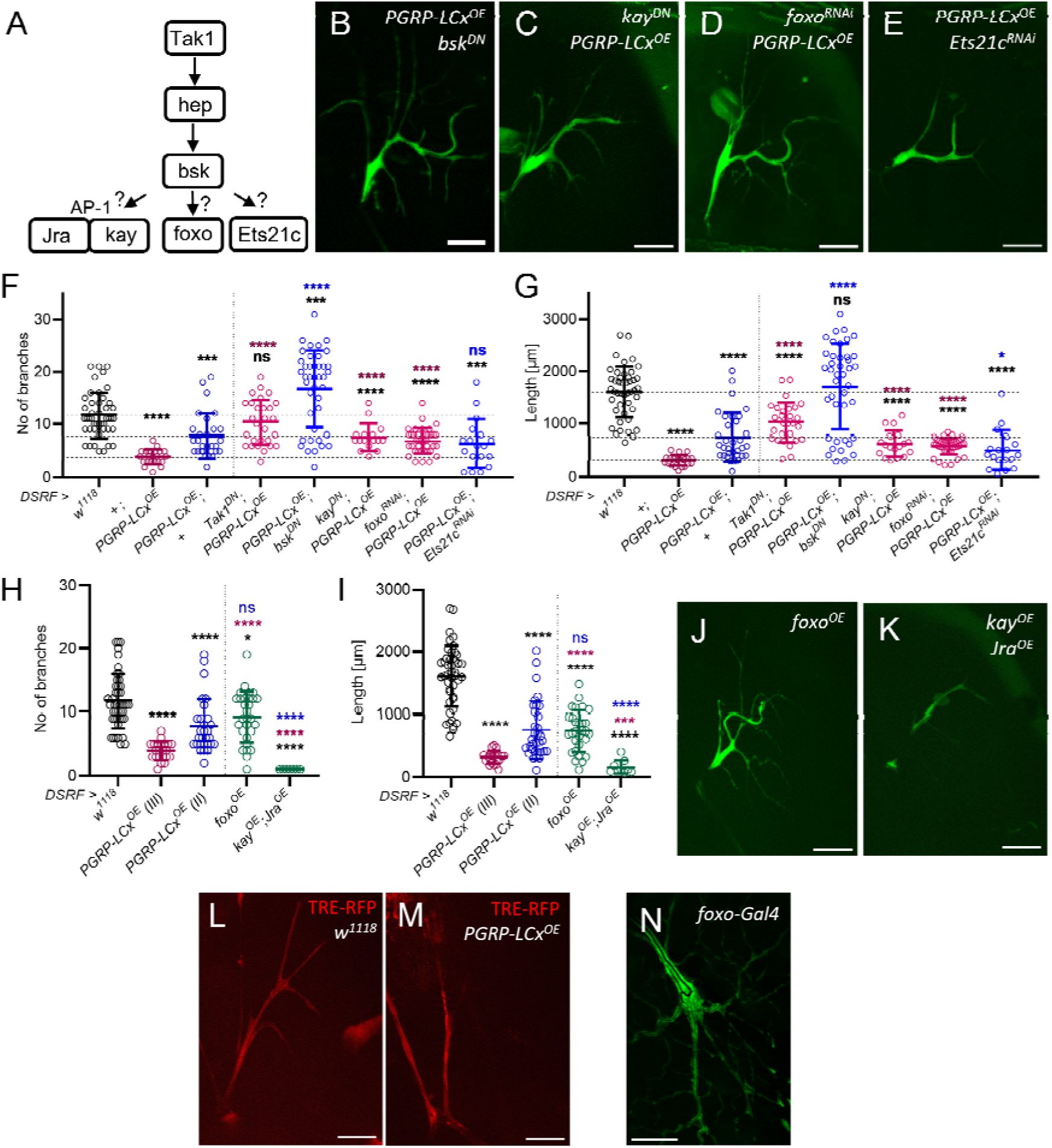
The TTC phenotype induced by PGRP-LCx is dependent on the transcription factors kay and foxo. (**A**) Schematic illustration of the JNK signaling pathway downstream of Tak1, which includes Ets21C, kay, and Jra (AP1). (**B–E**) *DSRF*-driven *PGRP-LCx^OE^* in TTCs was combined with the dominant-negative form of *Tak1, bsk* (B), and *kay* (C), or with RNAi targeting *foxo* (D) or *Ets21C* (E). (**F**, **G**) Measurement and quantification of the number (**F**) and length (**G**) of branches (n=16–45). **(H–K)** Measurement and quantification of the number (H) and length (I) of GFP expressing branches in control (*w^1118^*) and PGRP-LCx-expressing cells (n=7–45). TTCs overexpressing *foxo* (J, *foxo^OE^*), and *kay* and *Jra* (K, *kay^OE^* + *Jra^OE^*). (**L**, **M**). TRE-RFP expression in control (L) and PGRP-LCx-expressing TTCs (M). (**N**) *Foxo* promoter activity in TTCs (*foxo-Gal4 > UAS-GFP*). Data are expressed as the mean ± SD. Statistical significance was evaluated using Mann-Whitney-U test, * p < 0.05, *** p < 0.001, **** p < 0.0001, ns = not significant. The color of the asterisk indicates the corresponding comparison. Dashed lines represent the mean control value. Scale bar, 50 µm.

### Foxo controls TTC branching under normal conditions and under conditions of oxygen deprivation

A previous study reported a role for foxo in maintaining homeostasis of tracheal epithelial cells (Wagner et al. 2021). One of the main features that ensures full functionality of TTCs under changing conditions is the ability to adapt to local hypoxia to maintain the supply of oxygen to target tissues; therefore, we subjected wild-type larvae and larvae with foxo-RNAi in TTCs to mild hypoxia (5% O_2_) and measured their branching ability (Fig. 7). As expected, wild-type TTCs responded with increased branching (Fig. 7A, B, E). The *foxo^RNAi^*TTCs already had an increased number of branches under control conditions (21% O_2_; Fig. 7C–E), and the number of branches did not increase further after oxygen deprivation; this suggest that foxo is required for appropriate responses to hypoxic conditions (Fig. 7D, E).

**Fig. 7:**
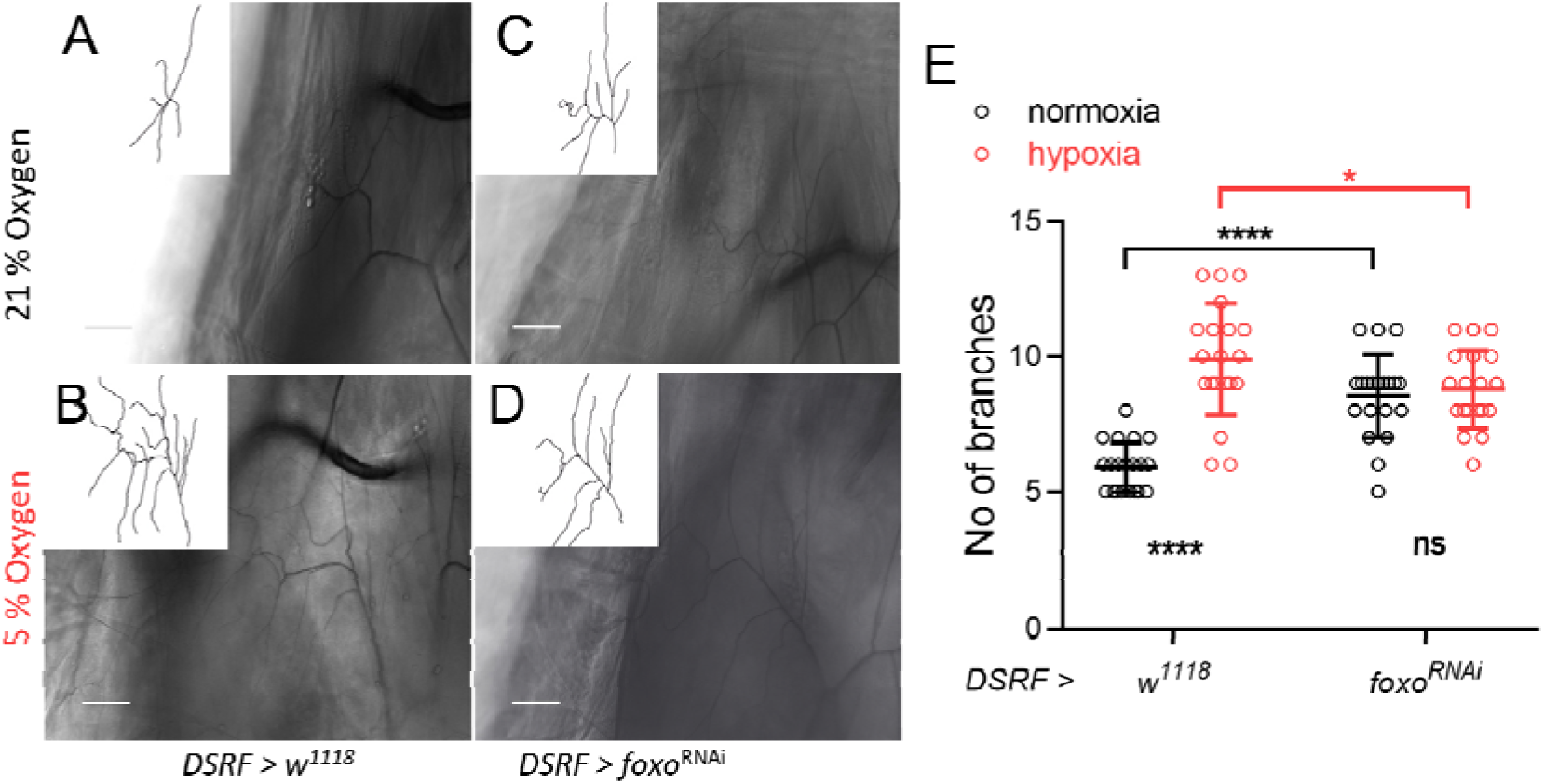
Targeted reduction of *foxo* expression in TTCs leads to hyperbranching. **(A, B)** Representative tracheal branching in control (A) and *DSRF*-driven *foxo^RNAi^*(B) cells under normoxic conditions. **(C, D)** Representative images showing tracheal branching in control (C) and *foxO^RNAi^*(D) cells under hypoxic conditions. **(E)** Quantification of branches in control and *DSRF > foxO^RNAi^* TTCs under normoxic (21%) and hypoxic (5%) conditions. Scale bar, 50 µm. n=21, Data are presented as the mean ± SD. Statistical significance was evaluated using Mann-Whitney-U t-test, * p < 0.05, **** p < 0.0001, ns = not significant.

In summary, the data suggest that TTCs differ from the other parts of the tracheal system in terms of immune pathway activation (Imd/JNK); this is a mechanism that circumvents cell death and prevents impairment of functionality (Fig. 8A). The Imd pathway receptor *PGRP-LCx* does not appear to be expressed in TTCs, indicating that no downstream signaling takes place (Fig. 8B). Ectopic expression of *PGRP-LCx* exclusively in these cells leads to JNK-mediated cell death; however, all downstream components must be present and functional. We have shown that depletion of the JNK-associated TFs AP-1 and foxo can rescue the PGRP-LCx-mediated phenotype. Both are present under physiological conditions, and their overexpression can induce a severe TTC phenotype. In addition, the universal transcription factor foxo, which is not only associated with JNK signaling, plays a role in branching of TTCs under normal and hypoxic conditions, indicating its importance for TTC homeostasis.

**Fig. 8:**
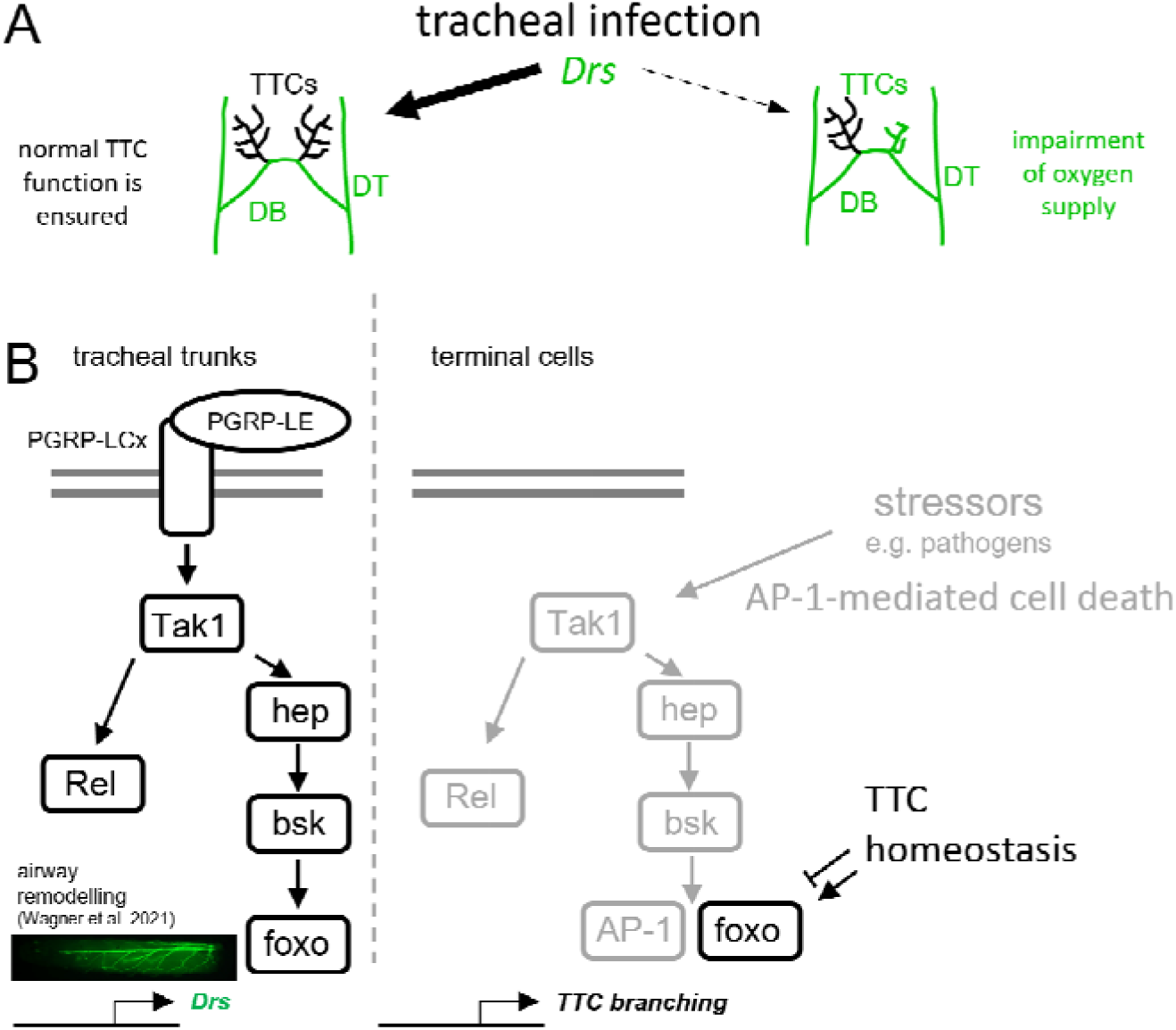
Schematic summarizing the JNK-mediated immune/stress response in the trachea and TTCs. (**A**) Tracheal infection leads to an immune response involving expression of antimicrobial peptides such as *Drosomycin* (*Drs*, green). In most cases, the immune response is restricted to the tracheal trunks and the TTCs are unaffected (bold arrow). In rare cases, TTCs express *Drs*, resulting in an impaired phenotype (dashed arrow). (**B**) Imd signaling in the main tracheal trunks is induced by peptidoglycan recognition receptors (PGRP)-LC and -LE. Downstream, the signaling branches into a Relish (Rel) and a JNK signaling pathway. Activation of the pathways mediates airway remodeling (*13*). However, activation in TTCs is avoided by the absence of PGRP-LC, even though all other JNK signaling pathway components are present. The pathway can be activated by external stressors, resulting in AP-1-mediated cell death. The transcription factor foxo, which is component not only of the JNK signaling pathway but also of the insulin signaling pathway, plays a role in TTC homeostasis and their ability to branch.

## DISCUSSION

Activation of the immune system is a double-edged sword in that although it fights infection, it can also damage the organism’s tissues (Ruff et al. 2020); therefore, appropriate regulation and restriction of the immune system is of prime importance. Tight regulation of the immune system is essential when delicate, sensitive tissues are involved, or when immune-relevant signaling pathways are also needed for other cellular functions. Tracheal TTCs appear to fall into both categories; they are susceptible due to their specific structure but require foxo signaling to maintain structural plasticity. Here, we found that TTCs, in contrast to the other epithelial cells in the trachea, barely respond to bacterial infection. Immune-privileged status might protect TTCs from damage caused by a strong immune response, a strategy that also protects other organ systems (Galea et al. 2007; Forrester et al. 2008; Hill et al. 2021). Here, we found that the entire tracheal epithelium expressed the transmembrane receptor PGRP-LC, but not TTCs. We propose that the lack of *PGRP-LCx* expression by TTCs is the reason for their lack of an immune response. This hypothesis is supported by the observation that targeted overexpression of *PGRP-LC* in TTCs induced an immune response, suggesting that all other pathway components are active in these cells. Since targeted overexpression of *PGRP-LC* leads to degeneration and, ultimately, apoptosis of these cells, this finding is particularly interesting. Targeted induction of apoptosis in TTCs by *hid* and *rpr* induced a similar, but more severe, phenotype; indeed, these larvae die after complete ablation of the cells. The weaker phenotype we observe in *PGRP-LCx* expressing TTCs could be also explained by a non-apoptotic function of Dcp-1. Dcp-1 encodes for an effector caspase during the apoptotic process but does also have non-apoptotic functions. Dcp-1 is involved in autophagy regulation and mitochondrial morphology (Hou et al. 2009; DeVorkin et al. 2014). In our case Dcp-1 is not only cleaved upon *PGRP-LCx* expression in the TTCs, but also in the tracheal epithelial cells. As described, *PGRP-LCx* expression in these cells does mediate tissue hyperplasia and renewal (Wagner et al. 2021). Furthermore, a role in inhibition of innate Toll signaling by cleavage of NF-κBs was reported (Wu et al. 2022).

Herein, we show that A) TTCs respond to a strong immune response by undergoing apoptosis, which leads to the death of the animal; and B) that immune activation is prevented by a lack of expression of the central receptor of the Imd pathway, PGRP-LCx. To better understand the first finding, we needed to elucidate the mechanism of apoptosis induction on the one hand and to identify mechanisms that still protect against infection on the other. Activation of the Imd pathway by ectopic expression of *PGRP-LCx* throughout the trachea leads to JNK-dependent meta- and hyperplasia of the trunk cells (Wagner et al. 2021). The JNK pathway, as well as activation of its canonical TFs, is associated with both proliferation and apoptosis. The intertwining of the Imd pathway with the JNK pathway occurs at the level of Tak1 and is a general organizational principle within these signaling pathways. This architecture means that strong activation of the Imd pathway in these cells also triggers the multifaceted JNK pathway. Downstream of JNK (*bsk* in *Drosophila*), we found that foxo and AP-1 are both necessary and sufficient for the apoptotic phenotype in TTCs. AP-1 is a classical TF that is activated by JNK signaling. The reason underlying the incomplete rescue is unknown but might be due to a specific role for foxo in this context.

We also demonstrated that depleting *foxo* rescues the apoptotic phenotype, at least in part. Although the effects triggered by *foxo* depletion or *foxo* overexpression were statistically significant, they were certainly not as strong as those triggered by AP-1. This difference in efficiency is because the tools used to manipulate foxo (RNAi and overexpression) are much weaker than those used to manipulate AP-1 signaling. For example, in the latter case, a dominant-negative form of *kay* and a fully active heterodimer of *Jra* and *kay* were available; in the case of foxo, the overexpression allele does not give rise to an activated version of foxo, and RNAi is generally less effective than dominant-negative alleles. In the tracheal cells of the dorsal trunk, and the primary and secondary branches, foxo plays a role in tracheal remodeling downstream of Imd pathway activation (Wagner et al. 2021). It seems reasonable then that foxo would also be activated in TTCs downstream of Imd signaling, which would in turn disrupt branching control and thus TTC functionality, as well as the ability to respond to environmental changes. Our central hypothesis derived from these observations is as follows: since foxo is essential for structural plasticity, and this property is critical for the survival of the organism, foxo should not be activated by other signals such as those from the Imd pathway. Foxo is activated not only by JNK signaling but also is a canonical member of the insulin signaling pathway. Linneweber and colleagues showed that insulin signaling, and especially the insulin receptor, is necessary for TTCs to respond (i.e., with reduced branching) to changing nutritional conditions such as low protein concentrations in food (Linneweber et al. 2014). Moreover, different insulin receptor alleles affect the size of TTCs (Burguete et al. 2019). Here, we showed that foxo is also part of this signaling pathway and that it is (presumably) required to translate different insulin signaling activities into structural changes in TTC branching. A very similar scenario for FoxO signaling is operative in endothelial cells (ECs). Insulin and VEGF signaling in human ECs targets FoxO to control its transcriptional activity directly via PI3K/Akt signaling (Abid et al. 2004; Abid et al. 2006).

An alternative explanation for the observed lack of an immune response in TTCs could be their maximal distance from the spiracles. In this scenario, a gradient of bacterial inducers along the tracheal system might be expected, resulting in a gradual decrease in immune activation from the spiracles toward the TTCs. However, this is not what we observed. In tracheae that displayed an immune response, the response was largely homogeneous along the entire length of the tracheal system, from the spiracles to the TTCs. Only at the transition to the TTCs did the immune response drop abruptly. This observation argues against the gradient hypothesis and suggests that TTCs are specifically excluded from the immune response.

Following our discussion of why it is sensible and necessary to omit this vital part of epithelial immunity from TTCs, we will focus on alternative strategies that operate to ensure that the cells are protected against infection. The strong immune response of the tracheal epithelium may be sufficient to protect the trachea, making an immune response in distant TTCs unnecessary. This strong and efficient immune response in the airway epithelia includes contact structures that connect the tracheal system with the outside world: the dorsal trunks and the primary and secondary branches (Wagner et al. 2008; Davis and Engstrom 2012; League and Hillyer 2016; Bossen et al. 2023). TTCs reside at the very ends of these systems, so potential pathogens would have to travel a long distance and escape powerful immune responses before reaching the TTCs. In addition, the diameters of 0.1–1 µm are so small that most bacteria will not gain access (Jarecki et al. 1999).

The mechanisms underlying immune privilege vary. The same is true for the mechanisms that dampen the immune response. Here, we found that the dampening of the immune response was almost complete (based on non-expression of the proximal receptor within the canonical signaling pathway controlling epithelial immunity). A similar strategy was reported in a subpopulation of intestinal epithelial cells, called enterocytes that do not express the Imd receptor *PGRP-LCx* (Fink et al. 2016). It was hypothesized that this prevents chronic MAMP-mediated activation of innate epithelial immune responses in these cells, which are constantly exposed to gut bacteria. Organs such as the liver, kidney, lymphoid organs, brain, and lung are immunotolerant (Amersfoort et al. 2022). Relevant immune pathways that induce apoptosis are dampened to different extents. In *Drosophila* epithelia, this dampening applies primarily to the Imd pathways because Toll signaling is not functional in these structures (Ferrandon et al. 1998; Wagner et al. 2008) Interestingly, the mammalian homolog of the Imd pathway, the TNF-α pathway, also invokes these protective strategies. This can be seen in ECs, which are the first cells to meet pathogens in the circulation and share some characteristics with TTCs. For example, they express several PRRs, as well as secrete pro-inflammatory cytokines to initiate immune responses (Mai et al. 2013) To prevent EC death and apoptosis, protective genes and negative feedback loops become active when immune pathways are triggered (Bach et al. 1997). For example, VEGF/VEGFR signaling is immunosuppressive (Yang et al. 2018) because it inhibits TNF-α induced apoptosis of ECs; VEGF inhibits secretion of TNF in a concentration-dependent manner (Spyridopoulos et al. 1997). For example, during angiogenesis and VEGFR activation, ECs express the anti-apoptotic gene *surviving*, which reduces caspase-3 activity and inhibits TNF-α-mediated apoptosis (O’Connor et al. 2000). The anti-apoptotic gene A20, when expressed in ECs, protects them from TNF-α-induced cell death, and inhibits inflammatory responses triggered by NF-κB activation (Daniel et al. 2004). Trabid, the equivalent of A20 in flies, is a negative regulator of the Imd signaling pathway (Fernando et al. 2014) TNF-α signaling, which induces death of various cell types, is a classical apoptotic pathway. TNF-α-induced apoptosis is highly regulated in ECs, which resist TNF-α-induced and physiological inflammatory apoptosis via TAK1 (Naito et al. 2019), which also regulates necroptosis and metastasis of ECs (Yang et al. 2019).

A recent study from our lab shows an inability of TTCs to respond to low oxygen level upon *Trabid* knockout, which supports our hypothesis that this signaling pathway converging onto foxo activation is necessary for this response (Bossen et al. 2025). The branching process in TTCs is comparable with that observed during angiogenesis, in which the outgrowth of new capillaries is regulated by VEGF/VEGFR signaling. Angiogenesis is influenced by events that induce survival or apoptosis in ECs; it can only be maintained by EC survival or inhibition of apoptosis mediated; for example, by growth factors such as VEGF/VEGFR and PI3K/Akt (Chavakis and Dimmeler 2002). Expression of the Ig-family member CD31 by ECs prevents cell death and renders them immune privileged (Cheung et al. 2015). When the TNF-α pathway is activated in ECs, CD31 is also activated; this in turn activates the Erk/Akt pathway and subsequent exclusion of FoxO3 from the nucleus, leading ultimately to inhibition of apoptosis *in vitro*. FoxO factors play a central role in controlling cell fate (i.e., apoptosis or proliferation). Constitutively active FOXO3a promotes EC apoptosis by downregulating protective factors, whereas dominant-negative FoxO protects ECs from apoptosis (Skurk et al. 2004). In line with this observation, loss of *mFoxO1* from murine ECs leads to uncontrolled overgrowth and hyperplasia. Moreover, FoxO controls endothelial quiescence by reducing glycolysis and mitochondrial respiration (Wilhelm et al. 2016). Active FoxO inhibits EC migration, whereas its silencing has the opposite effect (Potente et al. 2005). By coincidence, we discovered a similar role for foxo in *Drosophila* larval TTCs. We showed that foxo controls the branching of TTCs. Overexpression of *foxo* led to reduced branching, whereas *foxo* depletion increased TTC branching. We hypothesize that TTCs need to avoid activation of the Imd pathway because they depend on foxo to maintain full functionality in response to changing environmental conditions. The exact role of foxo in TTCs, and the signaling pathways involved, remain to be elucidated. Our study highlights the importance of *Drosophila* TTCs as a model for human ECs and thus may provide information that is useful for angiogenesis-related research.

The data presented herein demonstrate how immune privilege is necessary to maintain the functionality of specific cell types. In the case of TTCs, which should mount potent immune responses, the architecture and interconnectivity of the Imd (TNF-α) and JNK pathways mean that cell fate is inevitably linked to foxo. This is, however, incompatible with the role of foxo in controlling the structure and functionality of these cells under changing conditions. For these reasons, switching off the Imd pathway is the only solution to this dilemma.

## MATERIAL AND METHODS

### Fly lines and husbandry

Flies were raised on corn meal medium at 25°C. Tracheal humoral immune responses were tracked using *Drs-GFP* (BDSC_55707), which allows measurement of the GFP-tagged promotor activity of the antimicrobial peptide gene *Drosomycin*. Similar to this, we employed GFP reporters for Defensin, Attacin, Metchnikowin and Diptericin (Tzou et al. 2000). The expression pattern of PGRP-LCx was analyzed by crossing *PGRP-LCx-Gal4* (BDSC_77776) flies with *UAS-GFP* (BDSC_52262) flies. Tracheal driver lines *ppk4-Gal4* (Liu et al. 2003) and *btl-Gal4; UAS-GFP* (*tub-Gal80ts*) (Leptin Group, EMBL Heidelberg), or TTC driver line *UAS-GFP; DSRF-Gal4* (Gervais and Casanova 2011)), were crossed to the corresponding responder lines *UAS-PGRP-LCx(III)* (Kathryn Anderson, New York), *UAS-PGRP-LC(II)* (BDSC_30918), *UAS-PGRP-LE* (*Dipt.-lacZ,UAS-Flag-PGRP-LE/CyO*, Shoichiro Kurata, Sendai), *UAS-Rel68* (BDSC_55777), *UAS-Tak1* (BDSC_58810), *UAS-hid;rpr* (*UAS-hid; rpr;* Christian Wegener, Würzburg), *UAS-bsk^OE^* (BDSC_9310), TRE-RFP attP40 (Chatterjee & Bohmann 2012), *UAS-bsk^DN^* (BDSC_44801), *UAS-kay^DN^*(BDSC_7214), *UAS-Ets21C^RNAi^* (BDSC_39069), *UAS-foxo^RNAi^* (BDSC_27656), *UAS-kay^OE^* (BDSC_7213), and *UAS-Jra^OE^* (BDSC_7216), or *w^1118^* (for control crossing, BDSC_5905).

### Infection experiments

Natural infection of larvae was performed as described (Tzou et al. 2000; Akhouayri et al. 2011). The gram-negative bacterium *Pectobacterium carotovorum* (*Erwinia carotovora*, Ecc-15, 2141) was cultured overnight at 30°C in LB broth. The bacteria were pelleted by centrifugation for 20 min at 3200 × g, resuspended in PBS, and absorbance at OD_600_ measured. Next, 200 µl of the bacteria solution (OD_600_ = 160) was dropped into a vial containing developing 2^nd^ or early 3^rd^ instar larvae (3 days after egg laying) in standard cornmeal medium. After 24 h, the 3^rd^ instar larvae were used for microscopy. The larvae were heat killed in a drop of glycerol at 70°C for 20–25 s and arranged on a microscopy slide with the dorsal side facing up. GFP expression in the tracheal DB and TTC between the dorsal trunks was analyzed.

### Branching analysis

The 3^rd^ instar larvae were washed in PBS and deposited on a slide in a drop of glycerol. The larvae were heat killed at 70°C for 20–25 s and arranged dorsal side up before microscopy. Images of the TTCs were taken using the GFP channel (at 10× or 20× magnification). The right TTC from the 3^rd^ dorsal segment was chosen for analysis. Images were taken by an Axio Imager.Z1 (Zeiss, Munich, Germany) with ApoTome in Z-stack mode to capture all TTC branches. Measurement of the number and length of TTC branches was undertaken using the ImageJ plugin NeuronJ (Meijering et al. 2004). The branches were measured, as described previously (Jones and Metzstein 2011).

### Immunohistochemistry and microscopy

For microscopic and immunohistochemical analyses of intestinal TTCs, the gut of 3rd instar larvae (along with the connected TTCs) was dissected. As an alternative, a fillet dissection for the cuticle-attached TTCs was used for microscopy. The tissue was dissected in PBS, fixed with 4% PFA, and blocked with 5% normal goat serum (Sigma-Aldrich, Munich, Germany) before overnight incubation at 4°C with an anti-GFP antibody (1:200, DSHB, Iowa City, USA). Cells were also stained with anti-Dcp-1 (1:200, Cell Signaling Technology, Danvers, USA), anti-pJNK (Promega, V7931), and anti-Relish-C (DSHB, 21F3). Images were taken with an Axio Imager.Z1 (Zeiss, Munich, Germany) with ApoTome in the Z-stack mode.

### Hypoxia sensitivity assay

The 3^rd^ instar larvae were washed with PBS and placed into a new food vial with a scratched surface; 20 larvae were used per replicate. The vials were incubated for at least 10 min until all larvae were buried in the food and then deposited in a sealed desiccator. Nitrogen gas was introduced until the O_2_ concentration reached 2–5%. This O_2_ level was maintained for 25 min. The number of larvae outside of the food was counted every 5 min.

### Measurement of epithelial thickness of the dorsal trunks

For temporal control using the TARGET system, crosses (with *btl-Gal4; UAS-GFP tub-Gal80ts*) were maintained at 18 °C (restrictive temperature) until larvae reached third instar stage. Larvae were then shifted to 29 °C (permissive temperature) for 24 h to induce transgene expression. Late L3 wandering larvae were washed in PBS to remove residual food and dissected in PBS. Z-stack DIC and GFP images were acquired using an Axio Imager Z1 microscope equipped with ApoTome (Zeiss, Munich, Germany). Epithelial thickness was quantified using the AxioVision SE64 Rel. 4.9 software (Zeiss, Munich, Germany). Measurements were performed in the 9th tracheal metamere.

## Acknowledgments

We would like to thank Britta Laubenstein and Christiane Sandberg for their excellent technical assistance.

## Author contributions

Conceptualization: JB, TR

Methodology: RR, JB

Investigation: JB, LZ, RR, LS, JH

Visualization: JB, LZ, RR, LS, JH

Funding acquisition: TR

Project administration: TR

Supervision: TR, JB

Writing – original draft: JB, TR

Writing – review & editing: TR, JB, RR

## Supplementary figures

**Fig. S1:**
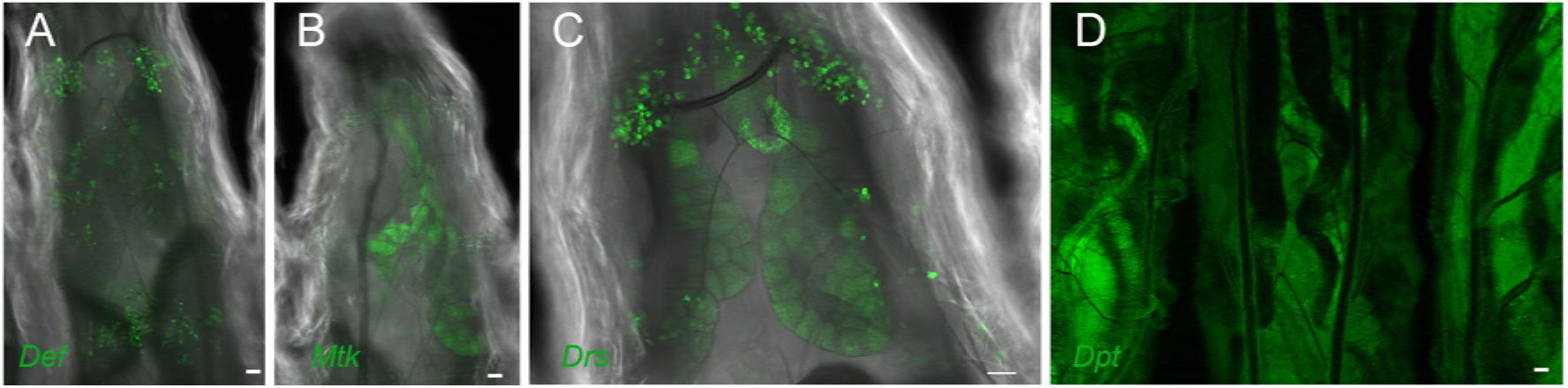
Drs expression in infected larvae. GFP reporter larvae were infected with *P. carotovorum* for 24 h, and GFP fluorescence was monitored. Drs expression was visible in different larval structures like hemocytes and fat body (A-D). Scale bar, 50 µm. Drs = Drosomycin, Def = Defensin, Mtk = Metchnikowin, Dpt = Diptericin.

**Fig. S2:**
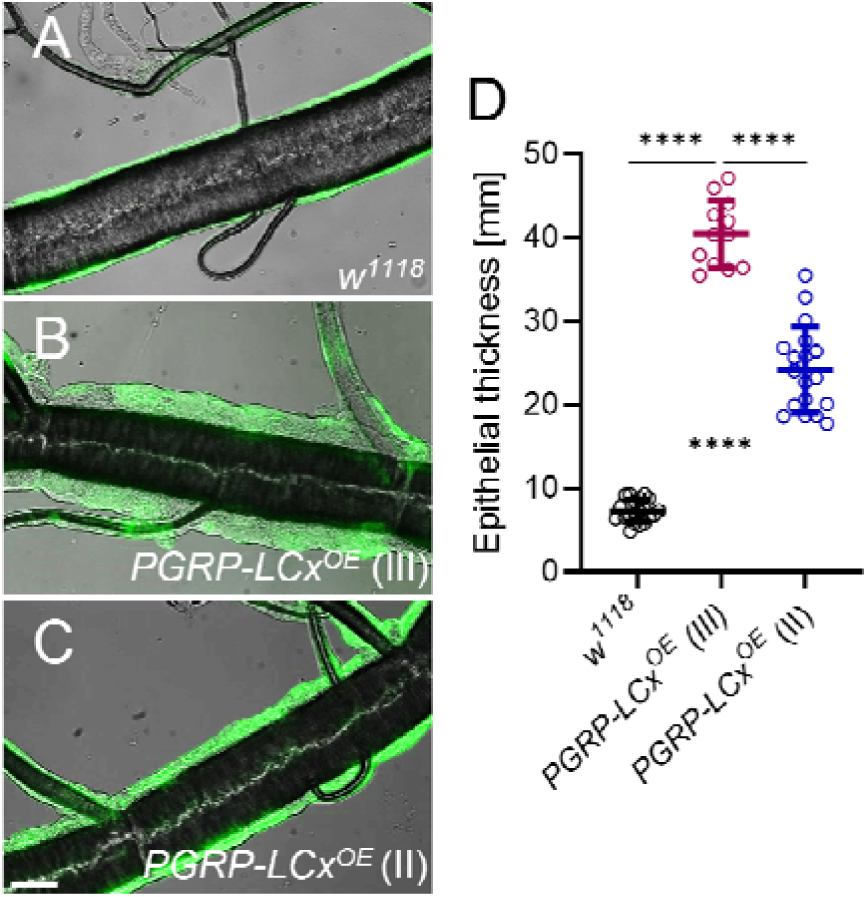
PGRP-LCx mediated epithelial thickening of the dorsal trunks. PGRP-LCx expression in the tracheal system was induced for 24 h in 3^rd^ instar larvae. Epithelial thickness of control larvae (A, *btl-Gal4, UAS-GFP; tub-Gal80ts > w^1118^*) were compared to PGRP-LCxOE larvae with PGRP-LCx on the third chromosome (III, B, *btl-Gal4, UAS-GFP; tub-Gal80ts > UAS-PGRP-LCx*) and on the second chromosome (II, C). Epithelial thickness was quantified (D). Scale bar, 50 µm. Data are presented as the mean ± SD. Statistical significance was evaluated using unpaired t-test, **** p < 0.0001.

**Fig. S3:**
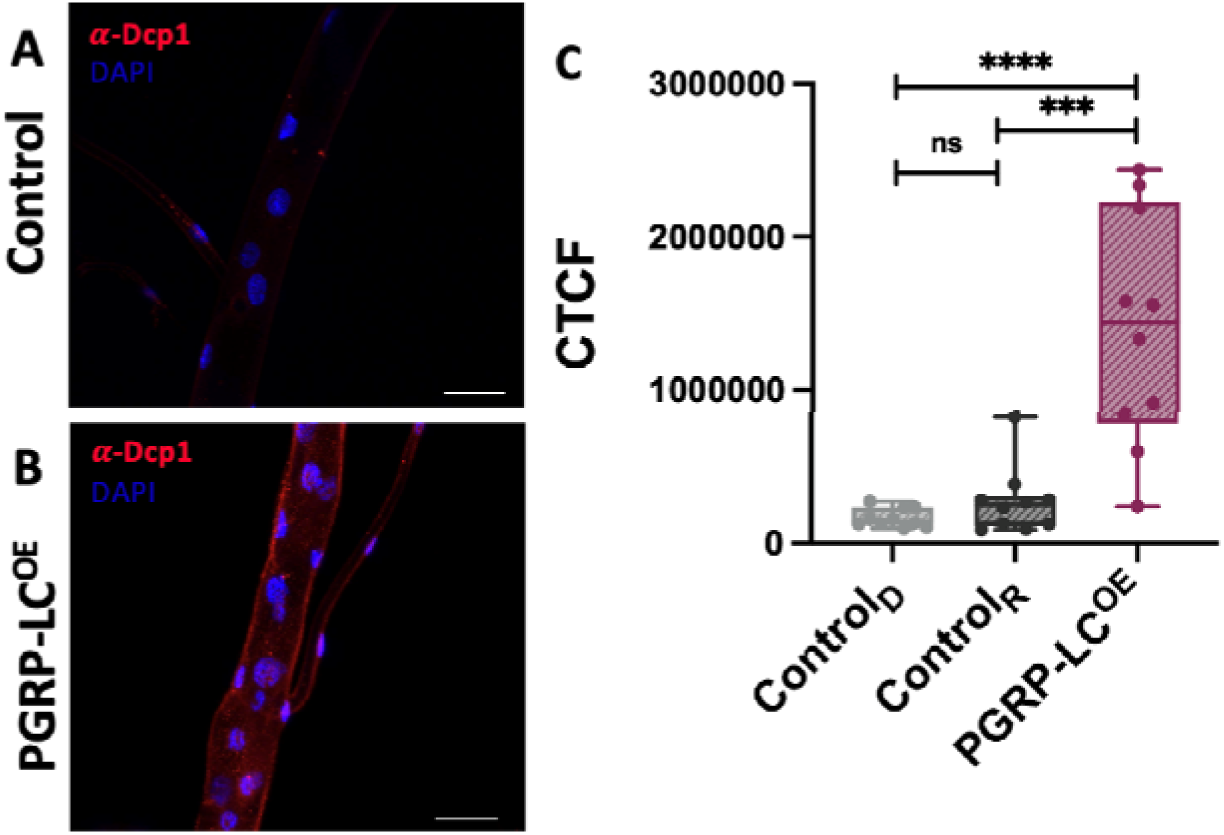
Increase in apoptotic signalling in PGRP-LC overexpression. The influence of PGRP-LC overexpression on apoptotic cells was investigated using -Dcp1 staining. For the analysis, the 8th tracheal metamere of the controls (**A**; *ppk4-Gal4 > w^1118^* (Control_D)_, *w^1118^ > UAS-PGRP-LC_x_* (Control_R_)) and PGRP-LE^OE^ (**B**; *ppk4-Gal4 > UAS-PGRP-LC_x_*) were used. (**C**) Quantification was performed by measuring the fluorescence intensity and calculating the CTCF (corrected total cell fluorescence) of Control_D_, Control_R_ and PGRP-LE^OE^. The statistical significance was evaluated using the Mann-Whitney test. ****= p < 0.0001, ***= p < 0.001, ns = not significant. Scale bar, 50 µm.

**Fig. S4:**
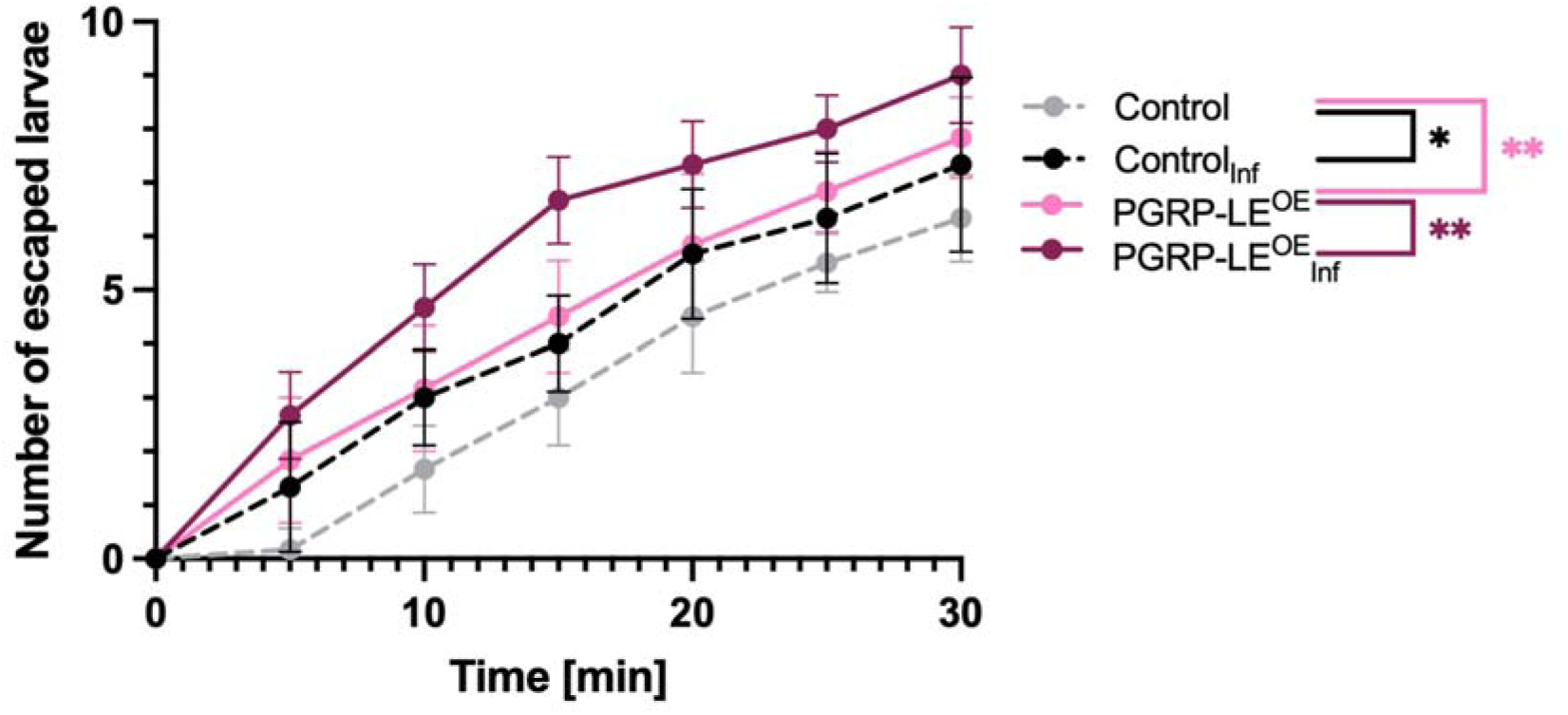
Overexpression of PGRP-LE and infection increased hypoxia sensitivity. The influence of PGRP-LE^OE^ and bacterial infection on hypoxia sensitivity was analyzed based on the escape response of 3^rd^ instar larvae exposed to 2.5 % OLJ. Control: *w^1118^ > UAS-PGRP-LE* (*Drs-GFP*), PGRP-LE overexpression: *Btl-Gal4;Gal80 > UAS-PGRP-LE (Drs-GFP*). Six independent replicates, each with 10 larvae of the control (grey), infected control (black), PGRP-LE^OE^ (pink) and infected PGRP-LE^OE^ (burgundy) were used for quantification. Values are presented as mean ± SEM. Statistical analysis was performed using two-way ANOVA. **= p < 0.01, *= p < 0.1.

